# A Modular, AI-assisted Digitization Toolkit for Resource-Constrained Herbaria: A Case Study from Zimbabwe

**DOI:** 10.64898/2026.07.20.739607

**Authors:** Langalenkosi Gatula, Ilja Bezrukov, Joseph Atemia, Christopher Chapano, Pioneer Gamundani, Clemence Zimudzi, Patience Chatukuta

## Abstract

Herbaria serve as invaluable spatio-temporal repositories of plant diversity information. Digitization of herbarium collections enhances the accessibility, discoverability, and long-term preservation of this important plant information, yet financial and infrastructural constraints often prevent herbaria in resource-constrained regions from digitizing their collections. Consequently, critical plant diversity data gaps remain due to underrepresentation of these collections in global biodiversity databases. Here, we describe an AI-assisted modular digitization toolkit specifically designed for herbaria operating under limited funding, developed and refined through our experience digitizing the crop wild relative (CWR) collection of the National Herbarium of Zimbabwe. The toolkit comprises three core components: (1) a portable, cost-effective photostation assembled from commodity parts, (2) a streamlined cascade workflow for systematic digital imaging, and (3) an AI-assisted data management pipeline for image quality control, label transcription, data analysis, and presentation. Compared to manual transcription and legacy optical character recognition approaches, AI-based transcription achieves lower time cost while maintaining high accuracy, and AI-driven data management delivers accessibility and reduced expenditure relative to conventional database infrastructure. The toolkit is designed to allow herbarium staff full autonomy over the digitization procedure, ensuring institutional ownership and the capacity for independent continuation beyond initial project support. By prioritizing affordability, modularity, and simplicity, this toolkit provides a replicable framework that may enable resource-constrained herbaria to locally generate high-quality scientific data for conservation and the sustainable utilization of plant genetic resources.

## 1. Introduction

There are 4035 herbaria across the world, and they together, house over 406 million specimens collected over the past four centuries and from varying geographies for scientific research, conservation, and education (1). These specimens document species distribution, phenological patterns and ecological changes over time (2,3). In southern Africa, over 90 herbaria collectively hold over four million specimens (4,5). The national herbarium of Zimbabwe alone holds over 500,000 specimens (6). Like many herbaria in the region, it operates under severe constraints in infrastructure and funding which hinder maintenance, documentation and development of its collection (7).

The digitization of physical herbarium specimens into high-quality digital images maximizes their scientific utility by broadening access, improving representation, preserving fragile specimens, and accelerating scientific discovery through remote access and computational analysis (8,9,10,11). Several digitization models have been developed across the world, such as the Smithsonian’s Rapid Capture System (12), DigiVol’s volunteer-assisted imaging framework (13), and the iDigBio Specimen Imaging Tools (14). Many of these solutions rely on either powerful high-end conveyor systems, advanced automation, or large volunteer bases (15,16,17). While digitization greatly enhances the scientific value of herbarium collections (24), it also raises important ethical considerations regarding traditional knowledge, data sovereignty, and equitable benefit-sharing, which must be carefully navigated in the digitization process (31). For resource-constrained herbaria, which are often located in biodiversity hotspots, these industrial-scale mass digitization models are unattainable due to high capital requirements, inadequate technical infrastructure, shortage of trained personnel, and unreliable internet connectivity (8,18,19).

To ensure equitable participation of resource-constrained herbaria in global biodiversity networks, a portable, low-cost, modular digitization toolkit tailored for the needs of herbaria and used in tandem with an affordable AI-assisted transcription workflow is required. Recent technological advances offer new opportunities to address this need. The proliferation of affordable digital cameras, cloud computing services and AI tools has lowered the technical and financial barriers to digitization (18,19,20). Concurrently, modular workflows and simplified data standards have reduced the expertise required for implementation (21,22,23).

We describe here a practical, modular digitization toolkit that we developed for the national herbarium of Zimbabwe. The toolkit is tailored for resource-constrained herbaria and is organized around three core components: (a) a portable and cost-effective imaging photostation built from commodity parts, (b) a cascade workflow for image capture and data mobilization, and (c) an AI-assisted data management pipeline for data analysis and presentation with minimal budget requirements.

## 2. Material and Methods

### 2.1 Photostation Design and Components

We built a custom compact, and portable single-user photostation from consumer-grade commodity components (Table S1, Supplementary File). The photostation was designed for affordability, ease of assembly, low maintenance, and minimal physical interaction with the camera. At the same time, it is sufficient to provide high-resolution images suitable for detailed morphological analysis.

We adapted a design from Davis *et al*. (22), and built the photostation around a modular and portable camera rig and supporting equipment. The complete setup occupies about 1 m^3^ of table space and can be assembled in less than four hours by four people with basic technical skills.

At the core of the setup is a rig assembled from pre-cut aluminum extrusions that offers a sturdy and adjustable frame for mounting the camera and lighting system (Vention GmbH [https://vention.io]). We used a mirrorless interchangeable lens camera with 24MP (Canon EOS R10), equipped with a 35mm F1.8 macro lens (Canon RF 35mm 1.8 MACRO IS) to capture high-resolution images. Consistent lighting was achieved using an LED panel (Kaiser PL240 Vario), which ensured specimens were evenly lit during photography. To manage specimen identification and traceability, we used a barcode scanner (Honeywell 1470G) along with a thermal label printer (Seiko) and compatible label rolls to generate and read unique barcodes for each specimen. Image files were saved directly onto an SD card and a local server (Dell OptiPlex 5000), which was additionally utilized for data processing. A 1TB portable SSD (Samsung T7 Shield) housed within a RAID enclosure (ICY BOX) served as redundant local data storage. For color and size standardization, each specimen image included a color control patch (KODAK) with a color reference chart and a cm/mm ruler, facilitating color calibration and size measurement across the digitized collection. A monitor, keyboard, and mouse completed the setup. The photostation was successfully assembled in half a day without special tools by personnel without technical training.

### 2.2 Cascade Workflow for Image Capture

The systematic 8-step digitization and data management process (Figure 1) commences with the development of a digital inventory of specimens. We prioritized type specimens, which serve as the official, permanent reference for a species’ name and scientific description. We focused specifically on crop wild relative taxa, which are a small subset of the herbarium collection. Binomial names of species were cross-referenced against the Global Biodiversity Information Facility (40) taxonomic backbone and Plants of The World Online (POWO) to detect misspellings or outdated nomenclature (51).

**Figure 1.**
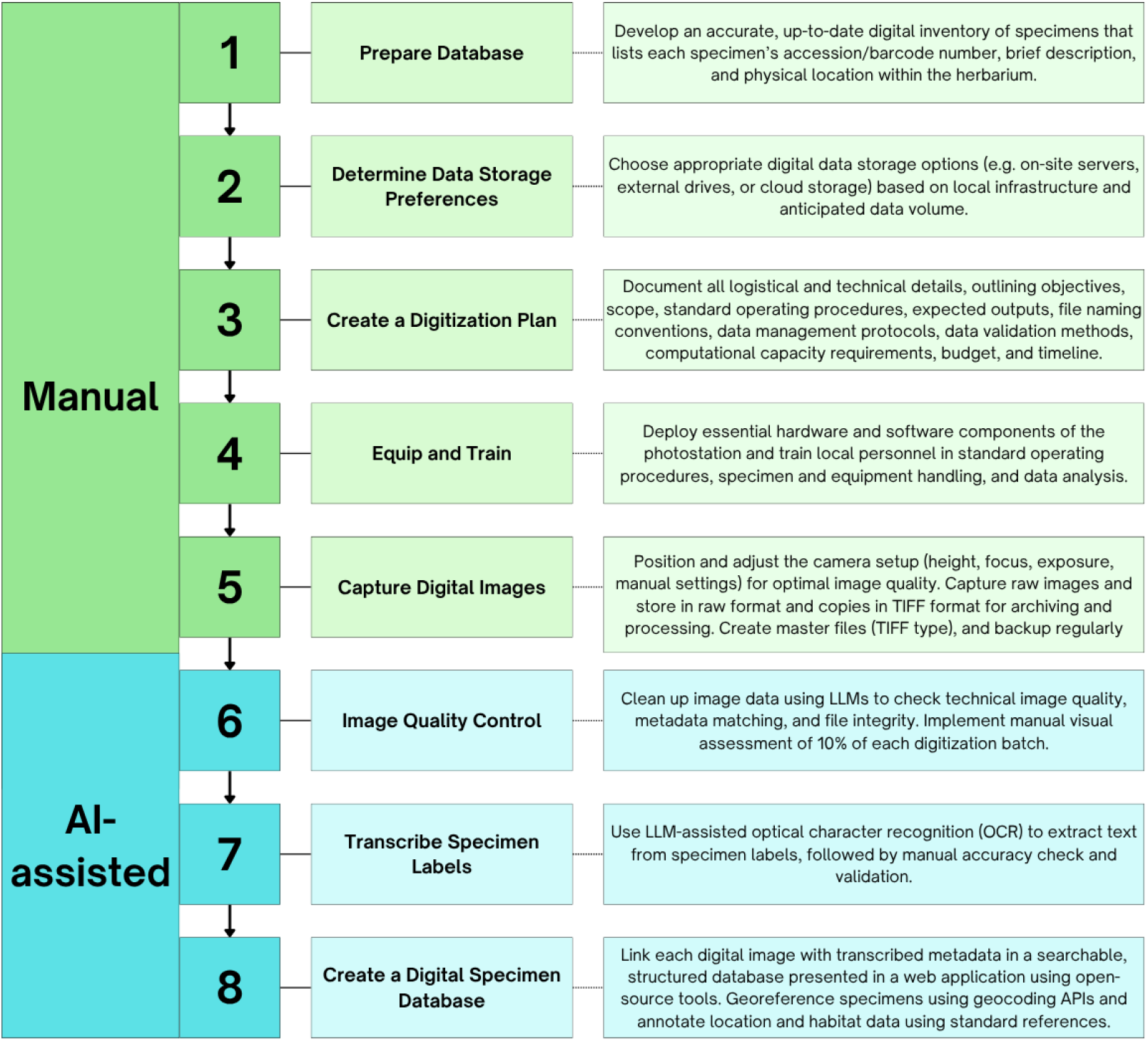
The 8-step cascade workflow for AI-assisted herbarium specimen digitization using a modular, portable photostation. The manual process starts with preparing the database, through creating a digitization plan, training personnel and capturing the digital images. The AI-assisted component relies on LLMs and APIs for image quality control, label transcription, and searchable digital specimen database creation, with human oversight.

#### 2.2.1 Specimen Inventory, Quality Evaluation and Barcoding

Specimens were located within the Herbarium using the Genera Siphonogamarum ad Systema Englerianum Conscript (Dalla Torre and Harms, 1900-1907). Specimens were examined for plant material integrity, utility of specimen label information, species representativeness, and digitization feasibility. To provide machine-readable traceability between physical specimens and digital records (2, 19, 41), barcodes were generated and assigned to each specimen, and barcode labels were affixed to the respective specimen sheets. Associated metadata, including the specimen’s binomial name, barcode identifier, physical storage location within the herbarium, and a concise morphological description, were recorded in the specimen inventory, prioritizing type specimens. Based on local infrastructure and the anticipated data volume, we chose to store the data on the local server, external hard drive and cloud storage.

#### 2.2.2 Equipment Setup and Personnel Training

A digitization plan was formulated to describe the procedures for image acquisition, data analysis, and data management. The photostation and various software to facilitate digitization were installed (Figure 2), including Canon Digital Photo Professional 4 and Canon EOS Utility for image capturing and analysis, Seiko Smart Label Printer 620 application and Microsoft Word for barcode printing, and Microsoft Excel for logging and tracking digitization metadata. A custom standard operating procedure (SOP) for specimen handling, digitization and quality control was established and personnel were trained to follow the SOP.

**Figure 2.**
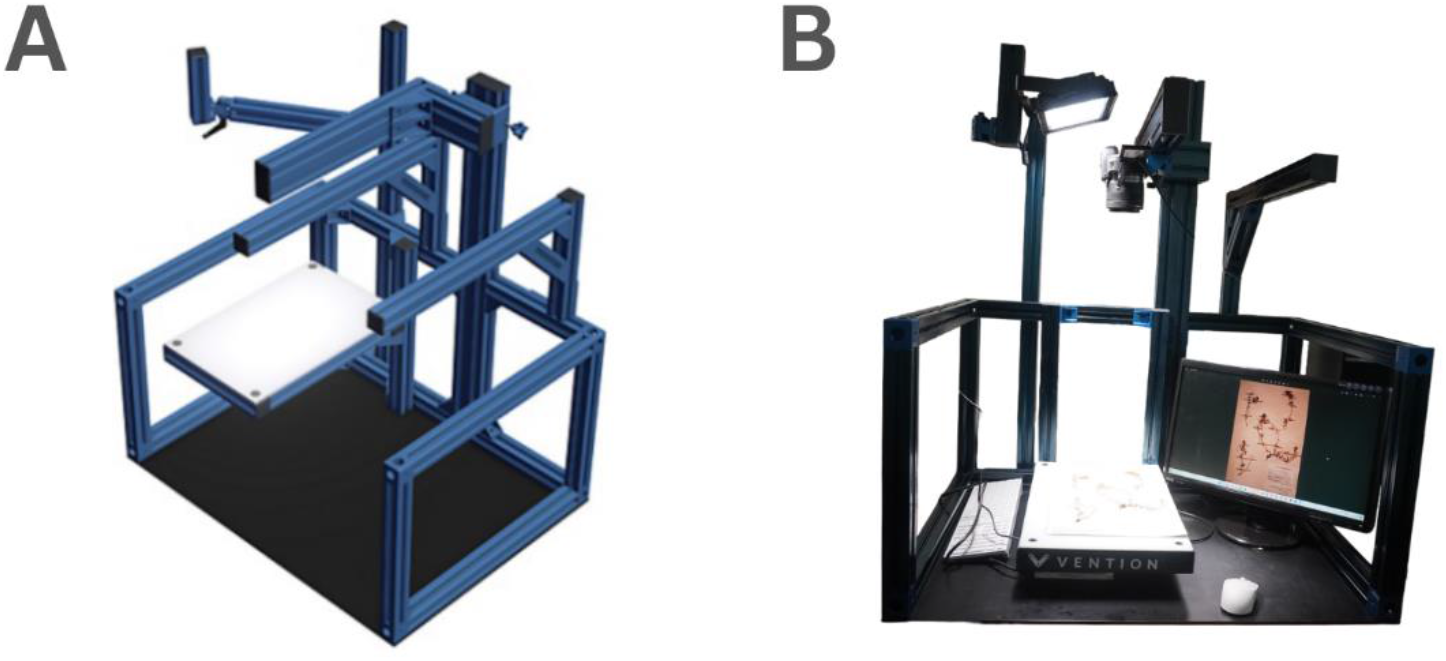
Photostation setup for the digitization of herbarium specimens at the Zimbabwe National Herbarium. (A). Modular rig to support photostation components, based on design by Davis *et al*. (23), with components manufactured by Vention GmbH (https://vention.io). (B) Assembled photostation composed of the modular rig and consumer-grade commodity components, including camera, LED light, specimen stage, and computer.

#### 2.2.3 Image Capture and Data Storage

Digital images were captured in accordance with our SOP (Figure S1, Supplementary File), and high-resolution master files were generated (Figure 3). Briefly, each specimen was placed on the photostation stage where it was within the camera’s range of focus. To ensure color fidelity and allow for post-capture white balance adjustments, a standard color control patch, the KODAK Color Control Patch, was utilized in every frame (36). The “Live View Shot” window on the Canon’s EOS Utility user interface was used alongside the LED lighting to precisely align the specimen within the frame and confirm optimal focus and exposure. Exposure was preset at the commencement of imaging. The image was captured via the Remote Shooting tool of Canon’s EOS Utility software. Each image was previewed for the following quality factors: camera focus, light distribution and camera’s flash intensity. Image data were automatically stored on the camera SD card and the server, and later uploaded to an external hard drive and cloud storage. Post-storage image quality was checked using the ChatGPT LLM, based on the ability to recognize a label from the picture, resulting in removal of misfired shots, blank frames, non-labelled frames and frames with extraneous material. All were checked manually to verify the LLM output. Specimen label data corresponding to the master files were transcribed and transcription output data were validated as outlined in Section 2.3.

**Figure 3.**
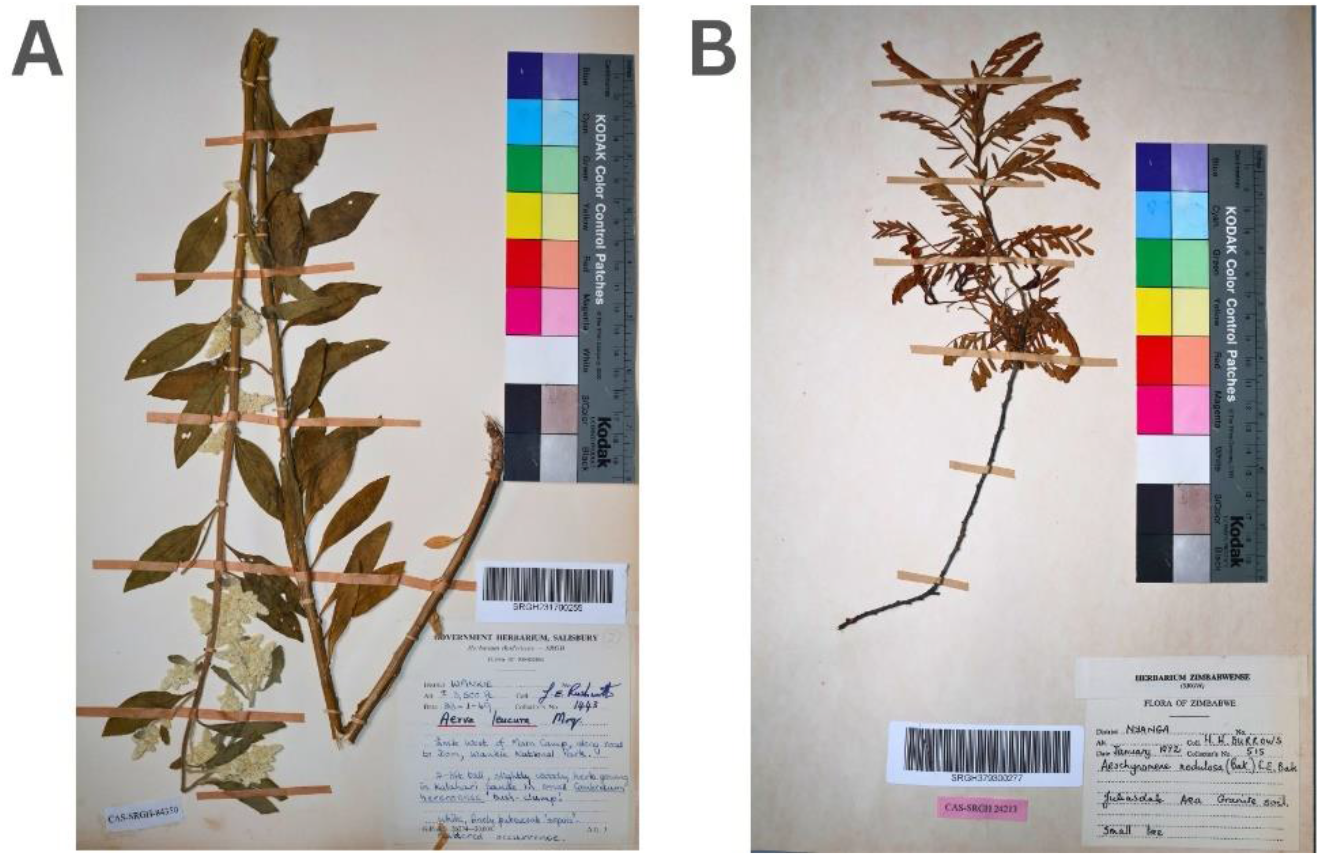
High-resolution digital images of specimens: (A) *Aerva leucura* Moq. and (B) *Aeschynomene nodulosa* (Bak.) E.E.Bak, housed at the National Herbarium of Zimbabwe and collected in 1969 and 1972 respectively. Both specimens have the label, color control chart and barcode attached. Images were acquired using the photostation at a resolution of 6,000 × 4,000 pixels (24 megapixels). Corresponding file sizes are 31.4 MB and 29.7 MB, respectively. Raw image files are stored in CR3 (Canon Raw 3) format and master files are stored in TIFF.

#### 2.2.4 Database Development

A searchable, structured database accessible through a web application was created to simplify access to the specimen images and transcription data as outlined in Section 2.3.

### 2.3 AI-assisted Specimen Label Transcription and Searchable Database Development

We extracted structured metadata from the labels of the digitized specimens and transferred these into a searchable database. The transcription and database development process entailed 4 steps: (a) transcription of handwritten or typewritten labels into the structured JSON format using a large language model (GPT-4o), (b) geocoding of each specimen based on the extracted metadata using the Google Maps geocoding API, (c) manual curation of the JSON data via a graphical user interface provided in a Jupyter notebook (Figure 4), and (d) making the data available through a web application (Figure 5). The transcription workflow with usage instructions is publicly available under an MIT license at github.com/ibebio/herb-transcribe.

**Figure 4.**
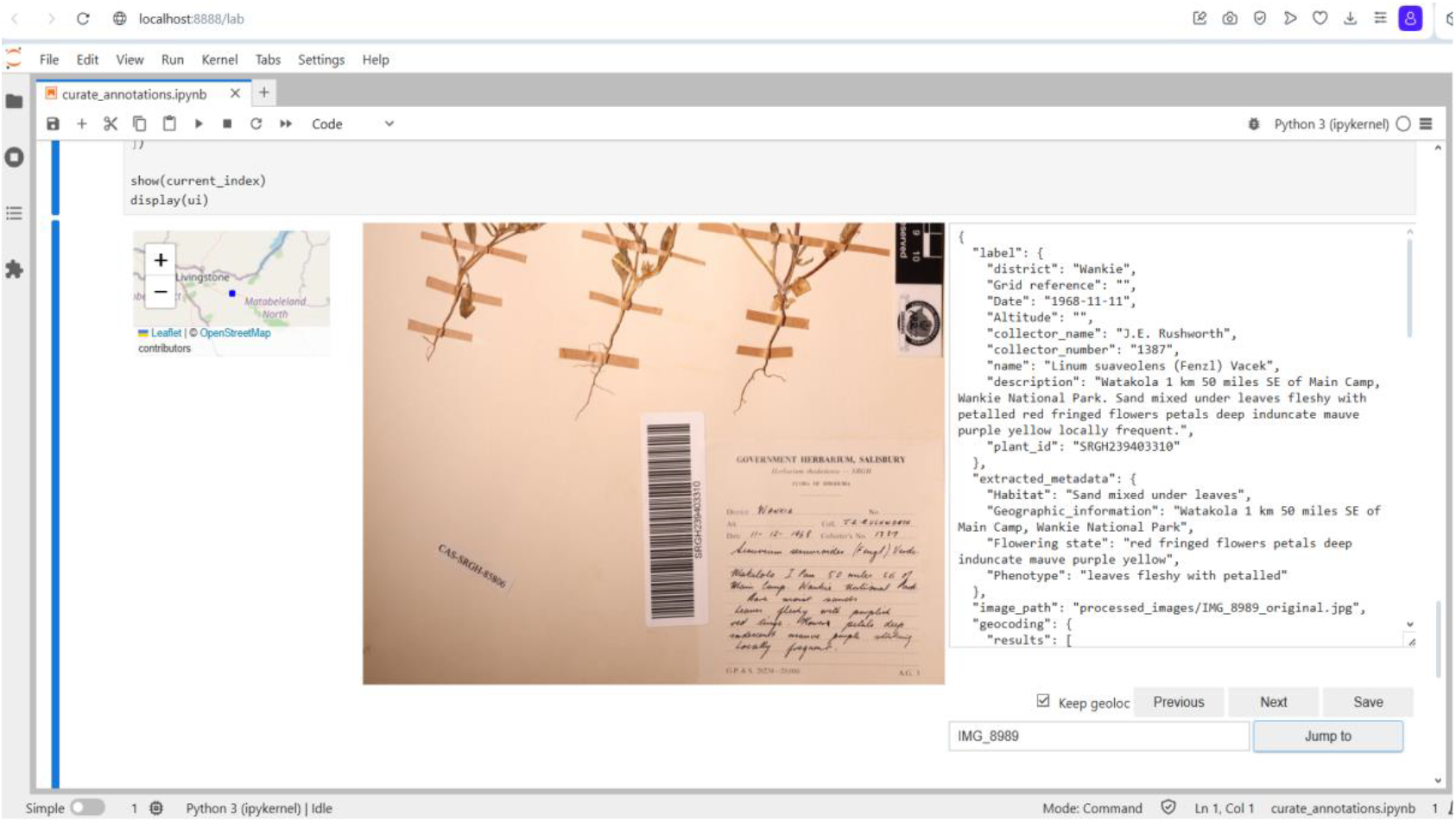
Jupyter notebook interface presentation of transcription and geocoding results. The specimen’s origin is shown in a map on the upper left-hand side. The image section with the label is shown in the middle. The GPT-4o-extracted data is on the right-hand side showing categories label transcription categories (district, grid reference, date, altitude, collector name, collector number, description, plant ID), metadata categories (habitat, geographic information, flowering state, phenotype) image path, geocoding and results.

**Figure 5.**
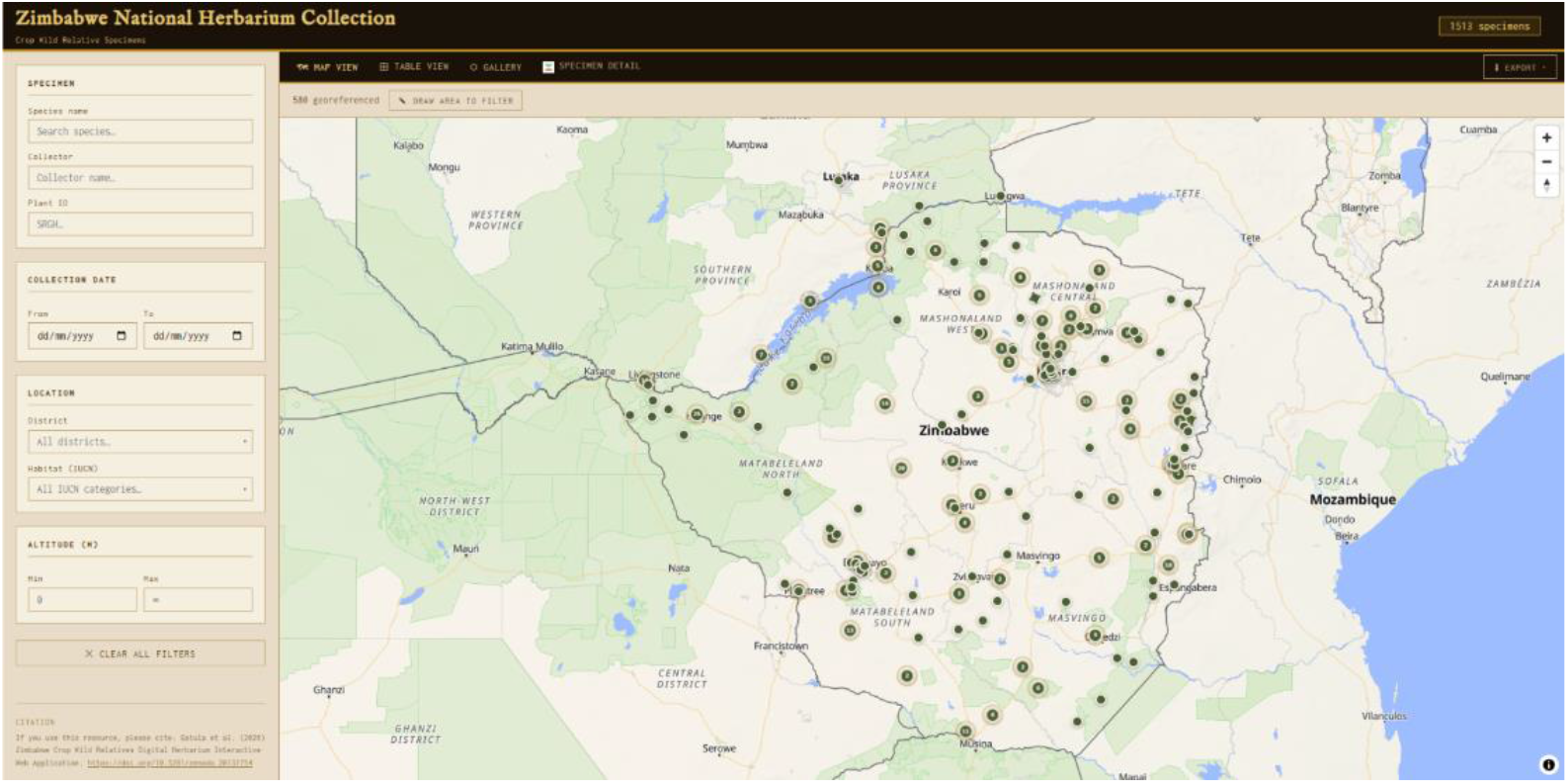
The web application (available at https://www.cwr.africanplantgenomics.org) presents the digitized specimen collection across four coordinated views: an interactive map displaying georeferenced specimens as navigable markers with hover popups and a draw-to- filter bounding-box tool; a sortable and filterable tabular view of the full collection; an image gallery groupable by district, habitat class, or collection year; and a specimen detail panel showing complete metadata alongside the digitized herbarium sheet photographs. Users can filter records by taxonomic name, collector, administrative district, IUCN habitat category, collection date range, altitude range, or a drawn geographic extent. Full-text search across taxonomic, locality, habitat, and descriptive fields is also supported. Filtered or complete datasets can be exported as CSV or a ZIP archive including images; exports require users to identify themselves and accept the data use terms via a short registration form with automated bot protection, and each export event is logged.

The web application integrates a relational database, a REST API, and an interactive browser-based interface to provide open access to the collection. Specimen records, geographic coordinates, and structured habitat classifications conforming to the IUCN Habitats Classification Scheme (IUCN, 2024) are stored in a PostgreSQL 17 (47, https://www.postgresql.org) relational database extended with PostGIS 3 (46, https://postgis.net), which provides native spatial data types, GIST-indexed geospatial queries, and bounding-box retrieval. Digitized herbarium sheet images are held on the server filesystem rather than in the database; a linked table stores only image metadata. Gallery thumbnails 400 pixels wide are generated once at ingest using Pillow v12.2 (43, https://pilow.readthedocs.io) and are served, together with the full-resolution masters, as static files by the nginx web server (52, https://nginx.org/). This keeps both Python and the database out of the request path for image delivery: the API is consulted only to resolve a specimen identifier to a filename, which it answers with an HTTP redirect to the corresponding static file. The backend REST API is implemented in FastAPI v0.133 (48, https://fastapi.tiangolo.com) with asynchronous database access via asyncpg v0.31 (44, https://github.com/MagicStack/asyncpg). The frontend is a React v19.2 (49) single-page application written in TypeScript v5.9 (53, https://www.typescriptlang.org/) and bundled with Vite v7.3 (50). Interactive map visualisation uses MapLibre GL JS v5.19 (45, https://maplibre.org), an open-source WebGL-based cartographic rendering library, and tabular data presentation is managed with TanStack React Table v8.21 (54, https://tanstack.com/table). Bulk export of specimen images is assembled client-side in the browser using JSZip v3.10.1 (55, https://stuk.github.io/jszip/). The application is available at https://www.cwr.africanplantgenomics.org (35); the source code is archived at github.com/AfricaPlantGenomics/digital-herbarium-web-app.

## 3. Results

### 3.1 Assessment of Image Quality

The digitization process prioritized minimal handling to reduce damage risk to specimens. On average, each specimen took about 5 minutes to digitize - this included scanning the barcode, capturing the image, and saving the digital file. Each acquired image had a resolution of 6,000 × 4,000 pixels (24 megapixels). Approximately 40 specimens were selected and captured per day and raw image files were stored in Canon Raw 3 (CR3) format. Images were processed for quality control through an AI-assisted data cleanup pipeline for removal of misfired shots, blank frames, non-labelled frames, frames with extraneous material, and redundant frames. Out of 4,437 images, 3,577 images passed quality control. An error rate of 19.38% (860 images) was calculated on redundancies (retakes). Redundancy decreased with time as proficiency with photostation operation improved. All errors were human errors and not associated with the equipment or SOP. A manual review confirmed that 100% of AI-detected errors were real and none were missed.

### 3.2 Data Handling and Storage

Data sustainability was maintained through the 3-2-1 Rule, where at least 3 copies of data were stored on two different media types, with one copy being stored off-site (26,41). All image files were saved directly to an SD card in the camera and to the local Dell OptiPlex server. The server facilitated review, backup, and management of files immediately after capture. Additional data backup was made onto the portable SSD and a cloud-based long-term archive.

For 3,846 unique processed images, total run time for automated AI transcription was 17h 01m 33s. A total of 2,497 images were selected for the database creation. Manual data validation of selected images via the graphical user interface (Figure 4) took 59 hours, therefore achieving a manual curation rate of 1.42min/image.

Transcription was conducted using OpenAI GPT-4o API at a marginal cost of €0.01/label and results were presented in a Jupyter notebook with the herbarium sheet image on the left side and a form containing GPT-4o-extracted metadata on the right side. Curators can quickly verify or correct each field in the notebook before approving the transcription. The curated metadata for 1,513 specimen records and their associated images were loaded onto the database and made accessible through the web application. The collection currently spans 117 years of collecting activity (1902–2019), documents the work of 550 individual collectors across 314 historical districts (reclassified to 54 modern administrative districts of Zimbabwe (28)), and is fully annotated with IUCN habitat classifications. Of the 1,513 specimens, 580 (38.3%) were successfully georeferenced; of these, 453 were via the Google Maps Geocoding API from locality text and 127 from Quarter Degree Square grid references.

### 3.3 Cost Implications

The entire photostation cost a total of €5,407.58, making it a highly affordable option for institutions working with limited budgets. Free to use open-source software was used where possible. Transcription of image labels of 2,746 images incurred total marginal costs of only €27.46.

## 4. Discussion

We prioritized type specimens and for the first phase, focused on the crop wild relative (CWR) complement, which is a small subset of the herbarium collection. Digitization of historical CWR specimens in Zimbabwe is aligned with the SADC Crop Wild Relatives Project (7,29), a regional effort to document CWR diversity in southern Africa. The CWR preselection also supported quality control by limiting scope to a manageable 2,497 specimens out of the over 500,000 specimens in the herbarium. The digitization of the full CWR type specimen complement of the herbarium allowed the development of a national checklist of CWRs emanating from the specimen inventory that provides a foundation for species prioritization for conservation and use (30).

By developing and testing a low-cost, herbarium specimen digitization toolkit at the National Herbarium of Zimbabwe, we have demonstrated that it is possible to produce high-quality digital images and manage data with AI tools effectively, without relying on expensive equipment or advanced technical skills. Compared to more advanced systems, this toolkit deliberately was designed to lack automation, because we prioritized affordability and ease of use. Importantly, the herbarium staff were able to take full ownership of the image collection process, allowing them to optimize and use the photostation independently of off-site technicians, with autonomy and according to context-specific priorities and timelines.

The main defining features of the toolkit are its modular architecture and its use of freely available LLMs and APIs, which allow herbaria to adopt components incrementally depending on available resources and evolving needs. It bridges the gap between the high-throughput systems of developed nations and the realities of herbaria in low-income countries, by prioritizing high quality data, portability, and affordability. The minimum viable system consists of image acquisition and cloud-based transcription, which reduces investment risk and enables resource-constrained herbaria to demonstrate value before committing to larger-scale efforts (27). Modularity supports customization to local contexts. Herbaria can substitute hardware components based on local availability and select software tools aligned with staff technical skills. Additional modules such as advanced geocoding or local collection management software can be added as capacity grows (21,22).

Our lockstep object-to-image model ensures only one imaging step after specimen inventory, which is more time efficient and gentle on the specimen compared to the object-to-data-to-image model commonly used by herbaria in the global South (34). The object-to-data-to-image model captures labels first and then records data from specimen labels. This model prioritizes pre-digitization curation; specimens may or may not subsequently be imaged, and are therefore precluded from specimen conservation consideration. Our model captures specimen images first, using the images as the basis for both label transcription and specimen conservation. Whereas the object-to-data-to-image model involves ‘before and after data’ specimen handling, our object-to-image model reduces specimen handling by ensuring only a single contact with the specimen.

The emphasis on pragmatic, and, where possible, open-source solutions (implied by the use of Python for the image quality control and transcription workflows, and cost-effective image transcription and web app development APIs) means the toolkit minimizes upfront investment while maximizing functionality and flexibility. Since the computationally heavy task of image transcription is performed using an external API, the manual curation can be performed on any laptop with a current operation system. While LLM services incur per-use costs, the efficiency gains can offset expenses for resource-limited institutions (19). Alternative open-weights LLMs such as Llama 3.2 vision, Qwen 3.5, and Gemma 4 may provide even more cost-efficient options as the technology matures.

The toolkit achieved a combined cost of €1.98 per image transcription, including the hardware cost. For institutions with tight budgets, a minimum viable system could be assembled for substantially less by eliminating the server and RAID enclosure, using existing computers for data processing, and employing free cloud-based storage solutions, potentially reducing the entry cost to approximately €2,500–€3,000 and transcription cost to €0.92-€1.10 per image. The per-specimen cost can decrease substantially with increasing transcription volume across a larger dataset, resulting in progressively lower marginal costs per specimen.

Assessment of the accuracy of label transcription by GPT-4o optical character recognition was challenging due to the inherent variability and non-standard nature of the label collection, which included handwritten and typewritten labels (25). The labels had non-standard formats, inconsistent punctuation, multilingual elements and variable handwriting styles. Although approximately 70% of the automated transcriptions required manual correction, the corrections varied in complexity. Corrections varied from standardizing ambiguous dates to adjusting collectors’ names to re-transcription of entire labels that were poorly legible. This made error rate calculation difficult, necessitating the use of a time-based metric to provide a more practical indicator of transcription efficiency. We measured the average time required per image for post-transcription review (1.45min/image), which proves the value of LLM-based transcription and yet underscores the need for expert oversight required for herbarium specimens.

Automated geocoding was constrained by the nature of the historical locality data, which frequently referenced obsolete place names, colonial administrative units, or imprecise vernacular localities that the geocoding API could not resolve [38]. Coordinates derived from Quarter Degree Square grid references carry an inherent spatial resolution of approximately 25–30 km, and the application’s visualization accounts for this by applying a small deterministic positional offset to grid-referenced markers to prevent visual overlap while preserving their approximate locality. Therefore, the derived coordinates should be treated as indicative rather than precise.

Our classification of all 1,513 specimen records against the IUCN Habitats Classification Scheme (37) at the point of data curation enables direct integration with Red List habitat assessments and facilitates queries that are not possible when habitat information is retained only as free-text. Similarly, standardizing raw altitude notation (feet, meters, ranges) to a single numeric field in meters preserves the original label data while enabling quantitative ecological comparisons across the collection. These enrichment steps add curatorial overhead but substantially increase the analytical value of the dataset beyond what the transcribed label text alone would support. Future development of the platform could include registration of the specimen records with the GBIF to maximize discoverability and support cross-collection species-level queries.

The web application and underlying database represent a significant step toward making an historically important but previously inaccessible southern African herbarium collection available for research. By providing spatially explicit, filterable, and exportable records through an open browser-based interface, the platform transforms a physical archive into a resource suitable for ecological, taxonomic, and conservation analyses that depend on occurrence and distributional data [39,40]. The use of an entirely open-source technology stack: PostgreSQL, PostGIS, FastAPI, React, MapLibre GL JS and nginx ensures that the application can be maintained, reproduced, and adapted without commercial licensing constraints. The only non-open-source dependency is the third-party bot-protection service used to gate bulk downloads, which is not required for the core browsing, search, and mapping functionality and can be substituted or omitted by adopting institutions.

The photostation described here addresses a fundamental need for practical, cost-effective digitization of herbarium specimens in resource-constrained regions where herbaria suffer from underfunding and infrastructure limitations, for the following reasons:

- Portability: All components are easily transportable and can be assembled/disassembled in under 4 hours. Components can be detached and relocated between rooms and buildings if need be.
- Modularity: Components are interchangeable with other makes/models, reducing reliance on specific vendors and offering flexibility in procurement.
- Ease of Use: Non-specialist staff can operate and maintain the photostation with minimal training, minimizing downtime and simplifying troubleshooting.
- Durability: Equipment is robust and easily replaceable, repairable and upgradeable.
- Space Efficiency: Entire setup fits within a compact footprint of approximately 1 m^3^ of table space, which is suitable for constrained working spaces without disturbing other operations.
- Data Security: Redundant storage on external drives and/or cloud ensures data safety and long-term stewardship. Barcodes act as unique identifiers to ensure permanent linking between the physical specimen and its digital representation. Local servers and external drives allow herbaria to maintain local control over data.
- Data Management: The interface, built as a Jupyter notebook, makes batch curation straightforward, with every validated record automatically marked for database ingestion. Label transcriptions are easily and rapidly done using LLM models, and specimen origin is easily geocoded with a suitable API based on locality data, eliminating the need for local software installation and leveraging cloud infrastructure (19). LLM-assisted transcription precludes intensive manual curation of transcriptions, saving time. AI-assisted coding tools enable accelerated web application development, enabling the rapid implementation of a full-stack geospatial platform that would otherwise have required extensive time and dedicated software engineering resources.
- Protection: The photostation can be easily covered for dust protection or during image capture, safeguarding both equipment and specimens.

While the toolkit provides a strong foundation, successful implementation at other institutions will depend on external factors such as consistent funding for consumables, personnel training, and dependable internet connectivity for API access, cloud storage and data sharing where applicable. While optical character recognition (OCR) and AI-based data extraction are continuously advancing, manual validation remains necessary to correct errors and ensure data accuracy (20). Reliable internet connectivity in resource-constrained contexts may be intermittent or costly, and therefore hybrid approaches that combine local processing with periodic cloud synchronization may need to be considered. Here, we used a commercial LLM service (OpenAI) with a low-cost implication; however, herbaria can explore open-weights model alternatives as they become available. Challenges include the initial cost of equipment and the need for trained personnel, as basic digital literacy and institutional commitment are required. These challenges can be mitigated through partnerships with international plant biodiversity networks to support capacity building (8).

Digital and open data publications raise important ethical, legal and social issues, particularly regarding traditional knowledge, access and benefit-sharing, and data sovereignty. Sensitive data, such as the location of endangered species, may require restricted access or redaction before publication (2). Herbaria can follow the Africa BioGenome Project’s guidance on implementing the Kunming–Montreal Global Biodiversity Framework (KMGBF) to help conserve biodiversity while promoting fair and locally beneficial use and sharing of biodiversity data (31).

Future work could involve developing detailed, open-source software packages specifically tailored for managing the digitized herbarium data generated by this toolkit, as well as exploring machine learning models for automated data extraction and analysis relevant to regional flora. Collaborative regional herbaria networks can coordinate digitization priorities and funding across multiple institutions, providing imaging services to smaller herbaria and facilitating knowledge exchange and capacity building. Future digitization workflows could also integrate specimen imaging and metadata with molecular data pipelines, enabling linking of morphological characteristics to genomic information (32,33,39,42).

## 5 Conclusion

The toolkit presented here demonstrates that affordable imaging hardware, streamlined workflows, and lightweight data management pipelines can be effectively deployed even in settings with limited infrastructure and funding. Our experience at the National Herbarium of Zimbabwe yields three key lessons: first, a modular photostation assembled from commodity components can produce high-resolution images suitable for research; second, large language models can dramatically reduce the cost and time required for label transcription, making data mobilization feasible at scale; and third, a full-stack open-source web application can transform a physical archive into a searchable, geospatially enabled resource accessible to the global research community. The successful digitization of over 1,500 crop wild relative specimens, spanning 117 years of collecting activity, demonstrates that resource-constrained herbaria can achieve meaningful and impactful digitization using the approaches described here. By enabling broader participation in global biodiversity research, this toolkit contributes to both scientific advancement and capacity building in under-represented regions, addressing geographic biases in biodiversity knowledge. Sustained commitment from national governments, funding agencies and the global biodiversity research community is needed to realize the full potential of herbarium digitization in resource-constrained contexts.

## Author Contributions

L.G. collected digital data, checked taxonomic nomenclature, and curated transcriptions. I.B. designed the photostation, and developed specimen label transcription and geocoding protocols. J.A. developed the web application. C.C. implemented photostation assembly and operation. C.C., C.Z, P.G, and P.C. conceptualized the study. All authors wrote the manuscript.

## Acknowledgments

L.G. funded by personal funds of Detlef Weigel. P.C. and J.A. funded by the Max Planck Society.

P.C. additionally funded by the Alexander von Humboldt Foundation. We are thankful for the support of the staff at the National Herbarium and Botanic Garden of Zimbabwe. We are also thankful to Martina Kolb for assistance with project logistics.

## Data Availability Statement

The searchable database of digital images and metadata of historical crop wild relative specimens from Zimbabwe can be accessed on the web database at www.cwr.africanplantgenomics.org. All specimen metadata, including geographic coordinates, are currently openly accessible; users are asked to identify themselves and accept the data use terms before downloading datasets. The Jupyter notebook and accompanying code for specimen label transcription can be found at github.com/ibebio/herb-transcribe. The web app source code can be found at github.com/AfricaPlantGenomics/digital-herbarium-web-app.

## Supplementary Information

### Supplementary File

Table S1: Photostation components with reference to specific models and costs accrued.

Figure S2: Standard Operating Procedure for the Zimbabwe National Herbarium Digitization Photostation.

## Notes

https://doi.org/10.5281/ZENODO.20137753

### Competing Interest Statement

The authors have declared no competing interest.

