## Supplementary Information for "A Modular, AI-assisted Digitization Toolkit for Resource-Constrained Herbaria: A Case Study from Zimbabwe"

**Supplementary File**

Table S1: Photostation components with reference to specific models and costs accrued.

*Model reflects specific model deployed. Cost (€) reflects cost in Euro as at 15.10.2024.

| Component | Role | Model | Picture | Cost (€) |
|---|---|---|---|---|
| **Camera Rig** | Provides stable platform for camera and light source mounting | Vention GmbH modular rig | 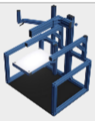 | 1.891,61 |
| **Camera** | Captures digital images of herbarium specimen | Canon EOS R10 | 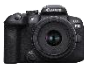 | 949,00 |
| **Lens** | Used in conjunction with camera to capture digital images | Canon RF 35mm F1.8 MACRO IS STM | 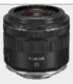 | 499,00 |
| **Light Source** | Ensures consistent and adequate lighting | Kaiser PL240 Vario LED Panel Light | 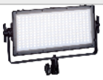 | 245,00 |
| **Barcode Scanner** | Reads specimen barcodes | Honeywell Hand Held Barcode Scanner 1470G:2D Black eMEA | 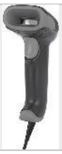 | 115,12 |
| **Barcode Printer** | Prints unique specimen barcodes | Seiko Instruments Smart Label Thermal Printer SLP620 | 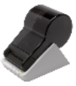 | 59,90 |
| **Barcode Labels** | Displays unique specimen barcodes | Seiko Instruments Address Labels SLP-2RLH 28mm × 89mm (5,200 labels) | 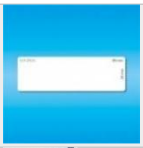 | 15,08 |
| **External SSD** | Stores data portably | Samsung Portable SSD T7 Shield 1TB MU-1T0R | 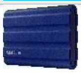 | 283,86 |
| **RAID Enclosure** | Protects external SSD | ICY BOX External RAID Enclosure | 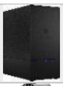 | 61,73 |
| **Server** | Provides local processing and storage capabilities | Dell OptiPlex 5000 Tower D32M001 | 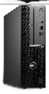 | 969,80 |
| **Colour Guide** | Enables colour calibration | KODAK Color Control Patch | 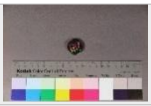 | 29,00 |
| **Accessories** | Monitor, Keyboard, Mouse, SD cards | | | 288,48 |
| **TOTAL** | | | | 5.407,58 |

Figure S2: Standard Operating Procedure for the Zimbabwe National Herbarium Digitisation Photostation

MAX PLANCK INSTITUTE FOR BIOLOGY TÜBINGEN

#### National Herbarium and Botanic Garden

### DIGITISATION PHOTOSTATION

#### STANDARD OPERATING PRODECURE

1. Switch on system plug, then switch on the CPU tower and the desktop monitor.
2. Connect camera, barcode scanner and barcode printer to CPU tower using the respective USB cords.
3. Switch on camera using the On toggle.
4. Switch on the barcode printer.
5. Open the program applications on the desktop monitor:
Canon Digital Professional 4, Smart Label Printer 620, Microsoft Excel data files

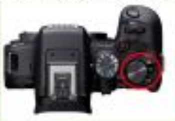

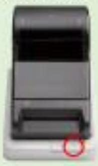

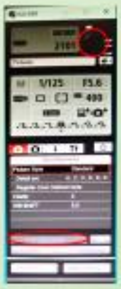

6. Select Remote Shooting in the EOS R10 application, then select Live View Shot in the pop-up window.
7. Switch on the LED light and observe feedback on the Live View Shot window.
8. Click on the barcode icon in the Smart Label Printer 620 application, enter the SRGH ID number/s, and click OK.

NB. SRGH ID number = SRGH+genus number+0+pictureID, e.g. SRGH035601785.
For genus with 3-digit code, add prefix 0.

9. Print barcode/s by clicking the print icon in the Smart Label Printer 620 application or by pressing ALT+P. Collect printed barcode/s.
10. Scan the barcode/s using the barcode scanner and then attach barcode/s to specimen/s.
11. Place colour control patch appropriately on specimen.
12. Place specimen on stage and align into correct frame guided by the Remote Live View window.
13. Capture specimen image by clicking on the round black button in the EOS R10 application.
14. Review the preview of the captured image.

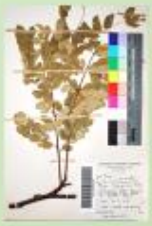

15. The image is automatically saved on the camera SD card and the daily folder on the Canon Digital Photo Professional 4 application. Ensure the daily folder is saved on This PC/Pictures/Zimbabwe/National Herbarium – Personal.
16. Remove the specimen and place it back in the folder.
17. Store colour control patch securely away from the light and ensure that it remains flat.
18. Close all applications on the desktop monitor.
19. Switch off scanner, LED light and camera. Shutdown computer and switch off monitor.
20. Switch off system plug.

Prepared by Langalenkosi Gatula and Patience Chatukuta.
Equipment kindly provided on loan from the Max Planck Institute for Biology Tübingen: https://www.bio.mpg.de
